# Evaluation of Microplate Handling Accuracy for Applying Robotic Arms in Laboratory Automation

**DOI:** 10.1101/2023.12.29.573685

**Authors:** Yoritaka Harazono, Haruko Shimono, Kikumi Hata, Toutai Mitsuyama, Takaaki Horinouchi

## Abstract

An inexpensive single-arm robot is widely utilized for recent laboratory automation solutions. The integration of a single-arm robot as a transfer system into a semi-automatic liquid dispenser without a transfer system can be realized as an inexpensive alternative to a fully automated liquid handling system. However, there has been no quantitative investigation of the positional accuracy of robot arms required to transfer microplates. In this study, we constructed a platform comprising aluminum frames and digital gauges to facilitate such measurements. We measured the position repeatability of a robot arm equipped with a custom-made finger by repeatedly transferring microplates. Further, the acceptable misalignment of plate transfer was evaluated by adding an artificial offset to the microplate position using this platform. The results of these experiments are expected to serve as benchmarks for the selection of robot arms for laboratory automation in biology. Furthermore, all information for replicating this device will be made publicly available, thereby allowing many researchers to collaborate and accumulate knowledge, hopefully contributing to advances in this field.

## 1. INTRODUCTION

Laboratory automation has significantly contributed to experimental science in terms of both quality and quantity. Moreover, recent research and development approaches, such as data-driven and AI-driven research, have proven to be powerful^1,2^, with laboratory automation being a crucial component. In the field of biology, commercially available automation systems are released from multiple vendors. However, the high cost of equipment installation hinders its widespread adoption. For instance, tabletop automated workstations commonly used in laboratory automation can cost tens of thousands of USD, and larger equipment can be several times more expensive^3^. Efforts are being made to reduce the system’s cost. In the case of liquid dispensing, which is among the most basic processes in biological experiment automation, semi-automatic dispensers such as the Opentrons’ OT-2 Liquid Handler^4^ and Integra Biosciences’ pipetting robot Assist^5^ have been introduced into the market at a more affordable price (a few thousand USD). In contrast, although fully automatic dispensers costing tens of thousands of USD are typically equipped with a gripper arm for sample loading and unloading, these semi-automatic dispensers lack a transport mechanism. Thus, they cannot handle sample transfers or connect with other devices without human intervention, rendering them limited-function automation devices.

An alternative to fully automated pipetting machines involves incorporating robotic arms for sample transfer in semi-automated pipetting systems. Robotic arms, with over six decades of application in the manufacturing sector for product transfer tasks^6^, have recently been applied in biotechnology. Robotic arms have been used for larger-scale systems, wherein fully automated pipetting machines and other large equipment have been integrated^7–10^ and broader application^11,12^. The advent of low-cost robotic arms by startups, priced significantly lower than traditional industrial robots^13,14^, along with educational robotic arms available for approximately 1000 USD^15,16^, has spurred their adoption in biological experiments. Several pioneering studies have already leveraged these affordable robots for specific tasks in biological experiments^17–19^. Considering their cost-effectiveness, we believe that robotic arms can enhance the functionality, such as sample transfer, of semi-automated pipetting machines, which otherwise have limited capabilities.

Position accuracy is crucial in robotic arm implementations for sample transfer. For instance, when moving labware such as microplates and tubes to a dispensing machine, precise placement at a designated location is essential. Misplacements can result in the loss of samples or reagents, potential equipment damage, or contamination. However, to the best of our knowledge, there exists a noticeable gap in the laboratory automation literature regarding the precise accuracy needed for successful transfers. Moreover, there is no quantitative data indicating the misalignment threshold that results in transfer failures. Manufacturers usually provide specifications for the repeatability of a robotic arm’s position (repeated accuracy of the robot arm’s position). Although the method to gauge this repeatability is detailed in ISO 9283^20^, it solely considers the robot’s approach to a specific location without holding any object. Numerous factors can influence this repeatability in practical scenarios. For instance, the position repeatability is affected when the end of the robot’s arms is under load derived from the labware and the end effectors attached to arms^21^. Extended operational durations can also introduce temperature-induced positional errors^22^. This lack of knowledge implies that the practical viability of a robot can only be assessed after its acquisition and during actual microplate transfers.

In this study, we implemented a commercially available robotic arm to simulate actual experimental conditions. A microplate and the corresponding end effector were attached to the arm to measure its positional repeatability. The hardware device developed for this purpose replicated a dispensing machine stage constructed from an aluminum frame. This stage included a microplate holder and dial gauges, which are used in the mechanical engineering field to measure spatial displacements^23,24^. Using this device, we moved the microplate to specific positions using a robotic arm and measured its position repeatability via dial gauges. Further, tests were conducted to analyze the effect of positional deviations on the probability of success of microplate transfers by deliberately offsetting the placement by a predetermined distance. The results of these experiments are expected to benefit users in the field of biotechnology when choosing a robotic arm to augment semi-automated dispensing machines. Furthermore, the proposed measurement device is cost-effective, easy to implement, and easy to operate. It has the potential to contribute to the accumulation of foundational data promoting the adoption of robotic arms in the automation of the biotechnology sector.

## 2. Materials and Methods

### 2.1 Code availability

The Python and Lua scripts used in this study are available at the GitHub repository: https://github.com/harazono/Evaluation_of_ScalaArm_Robot.

### 2.2 Creating in-house jig

The jigs used in this study were created in-house using a three-dimensional (3D) printer. The computer-aided design (CAD) data were designed using Fusion 360 Ver. 2.0.16009 x86_64 (Autodesk, Inc., CA, US). The created CAD data were uploaded to Supplemental **File S1**. The jigs were printed using a Raise3D Pro2 3D printer (Raise 3D Technologies, Inc., CA, US), and the filament used was a polylactic acid-based filament, RP-22-03 (Raise 3D Technologies, Inc., CA, US), with a diameter of 17.5 mm. We ordered the fabrication of a part of the jigs via an aluminum frame manufacturer (Misumi Group Inc., Tokyo, Japan). We purchased and used the designed jigs in the housing design software provided by the manufacturer, Misumi Frames^25^. The size of each aluminum frame is shown in Supplemental **File S2**.

### 2.3 Robot arm for plate transfer

We used Dobot M1 Pro (Shenzhen Yuejiang Technology Co., Ltd., Shenzhen, China)^26^ as a robot arm to conduct microplate transfer experiments. The end effector, an Electric Gripper (SERVO Type) DT-AC-PGE2-001 (Shenzhen Yuejiang Technology Co., Ltd.), was attached to the robot arm using an adapter DT-M1-FRG-001 (Shenzhen Yuejiang Technology Co., Ltd.). We attached fingers printed via a 3D printer (**Files S1**) to the gripper using 3 mm screws. Further, 1.0 mm thick silicon rubber (6-586-09, AS ONE Corp., Osaka, Japan) was adhered to the fingers as an anti-slip sheet. The robot was fixed to the experimental table using two clamps (PLW-200, Bakuma Industrial Co., Ltd., Niigata, Japan). The overview of the robot arm and end effector is shown in **Figures 1A and B**. In addition, we used DobotStudio Pro v2.6.0 (Shenzhen Yuejiang Technology Co., Ltd.) to program the movement sequence of the robot arm. The program was uploaded to the GitHub repository, as described in Section 2.1.

**Figure 1.**
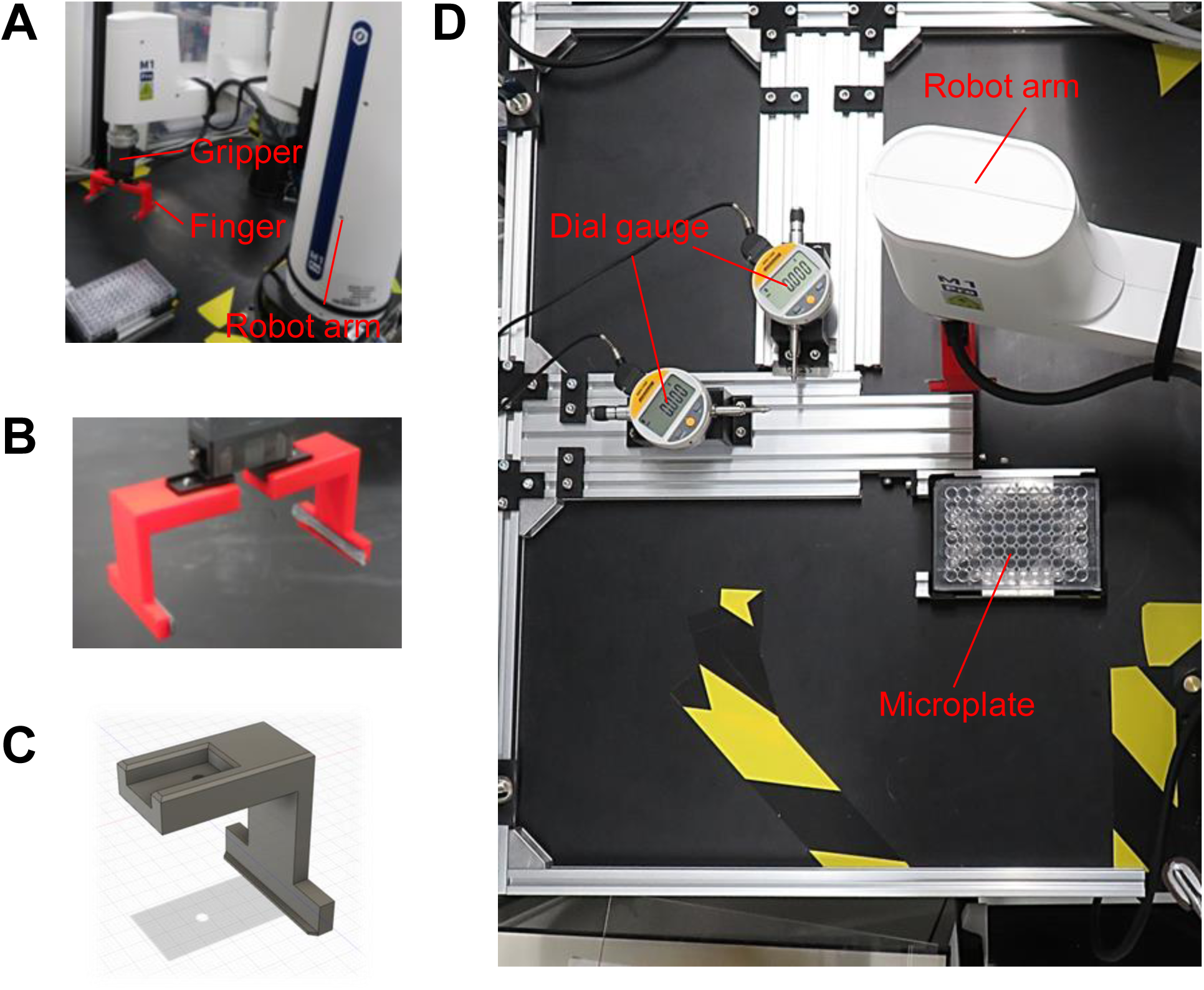
System for evaluating the microplate transfer. (A) Overview of the robot and end effector for transferring. (B) 3D-printed finger object attached to a gripper. (C) Screenshot of the 3D-printed finger object. (D) Bird’s eye view of the system.

### 2.4 Setup of stage for experiments

We constructed a stage where the necessary labware and measurement devices could be fixed within the range of motion of the robot to measure the position repeatability and acceptable error in microplate transfer using the robot arm. An overview of the stage is presented in **Figure 1D**. Detailed dimensions are presented in Supplemental **Figure S1**. First, we fixed the aluminum frame, which was cut to an appropriate length, to the experimental table (1200 mm × 750 mm, VQSA-DW-1200, Oriental Giken Inc., Tokyo, Japan) using clamps (CS-65, Ichinen Axess Co., Ltd., Osaka, Japan) (Supplemental **Figure S2**). Subsequently, jigs were placed to fix two digital gauges for measurement (DGN-125B, Ozaki MFG. Co., LTD., Tokyo, Japan) and a microplate holder. The digital gauges were set as parallel to the floor, and the long and short edge directions of a 96-well microplate (Corning, 3595, Corning Inc., NY, US) were set as x and y measurement axes.

Each digital gauge was connected to a computer for measurement (**Figure 2A**). Owing to the specifications of the digital gauges used, the request for data transmission was performed through serial communication using a USB cable and adapter (communication cable: KB-232C, input adapter: IF-21B, Ozaki MFG. Co., LTD., Tokyo, Japan), and BLE communication was used to receive the measured values.

**Figure 2.**
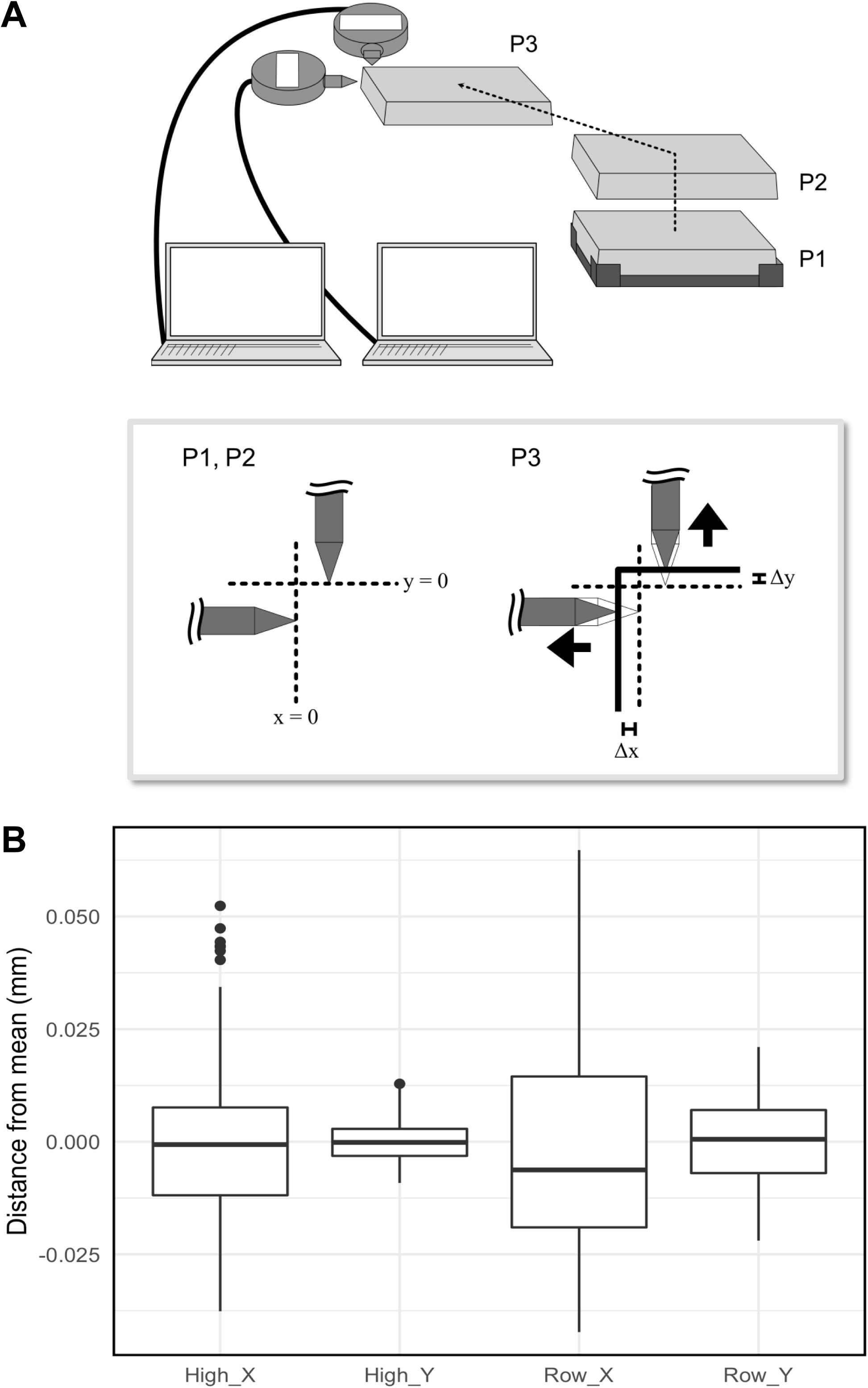
(A) Overview of the error measurement experiment in microplate transferring. First, the microplate is lifted (P1→P2), and then transferred to a designated location and pressed against the dial gauge’s probe (P2→P3). The amount the probe is pushed by the microplate (Δx, Δy) is recorded, and this process is repeated. (B) Measurement results of the error when repeatedly transferring the microplate (n = 125).

### 2.5 Experimental design for the evaluation of position repeatability of transfer microplates

We used a Dobot M1 Pro to transfer the microplate to a predetermined location and measured its displacement using the digital gauges to evaluate the position repeatability of the robot arm. This sequence of movement and measurement was iterated several times. A 96-well microplate was used for transfer. A measurement request was sent to the digital gauges at intervals of 2 s, and the measurement data were saved on a Google Spreadsheet. The operation sequence performed by the Dobot M1 Pro is presented in **Table 1**. The measurement values from the digital gauge were extracted for each repetition using the in-house script (repeatability_measurement.py) provided in our GitHub repository in Section 2.1. The extracted measurement values are presented in Supplemental **Table S1**. The code for this experiment was placed in the directory Evaluation_of_the_position_repeatability_of_transfer_microplates in our GitHub repository in Section 2.1.

**Table 1:** Operation sequence for evaluation of position repeatability of transfer microplates performed by Dobot M1 Pro.

| Step | Description |
| --- | --- |
| 1 | Open the end effector and move it to the top of the lid (as expressed P2 in Figure 2A). |
| 2 | Go down vertically from P1 to the height of the lid (P1). |
| 3 | Close the end effector and grip the lid. |
| 4 | Move to P2. |
| 5 | Move horizontally from P2 to the measurement position (P3). |
| 6 | Wait for 3 seconds to measure values of digital gauges. |
| 7 | Move to P2. |
| 8 | Move to P1. |
| 9 | Open the end effector and put the lid on. |
| 10 | Move to P2. |

### 2.6 Experimental design for the evaluation of magnitude of positional misalignment of transfer microplates

We evaluated the acceptable positional error that would result in the successful placement of the lid on a 96-well plate. We repeatedly opened and closed the lid by artificially introducing offset distances. Evaluations were conducted on the same stage as depicted in **Figure 1D**. The outline of the evaluation method is shown in **Figure 3A**. The operation sequence of the robot arm used for the measurements is presented in **Table 2**.

**Figure 3.**
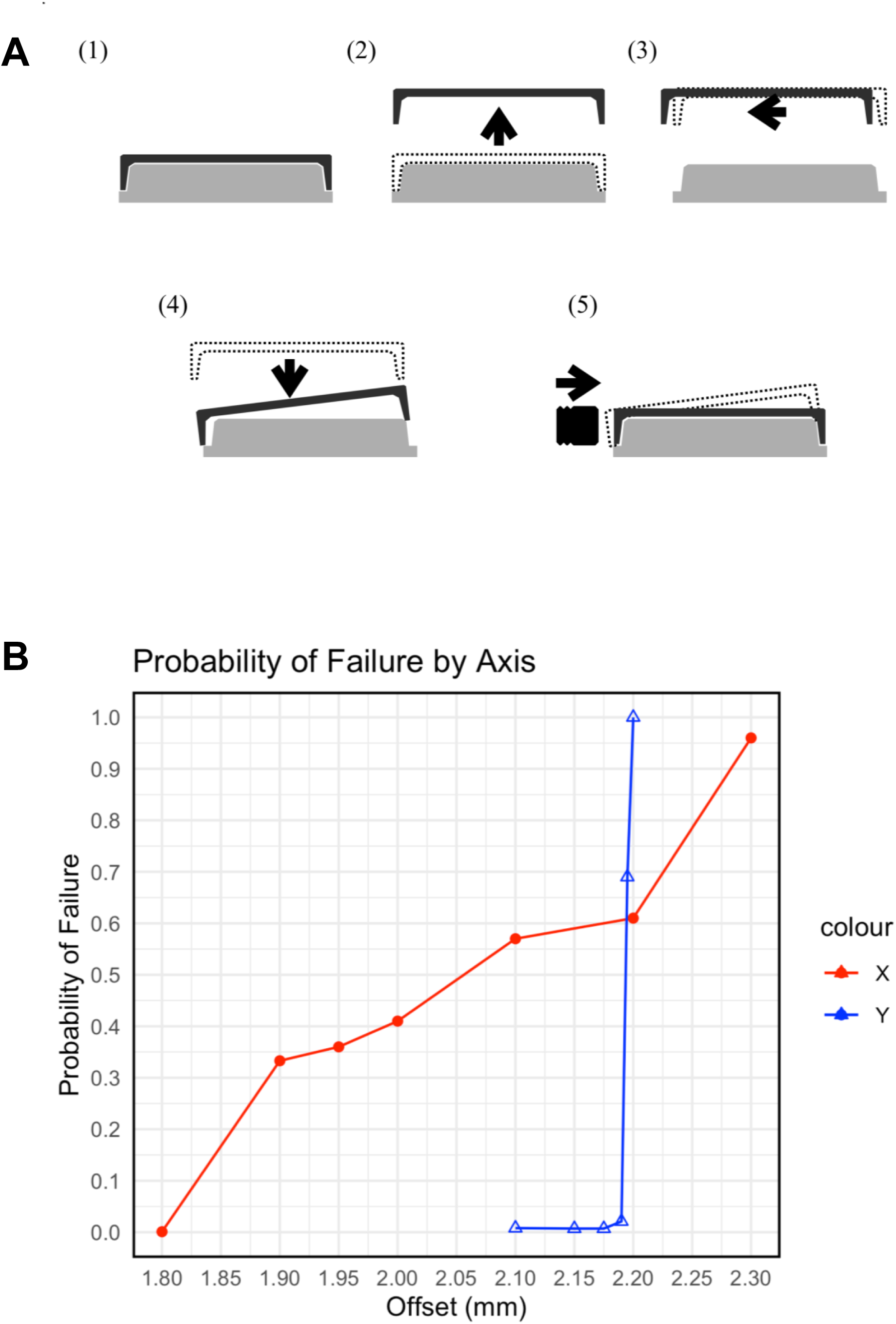
(A) Overview of the experiment to evaluate the success or failure of transfer when a positional deviation is introduced intentionally. First, the lid of the microplate is lifted (1→2), then moved a specified amount in either the x or y direction (2→3), the lid is lowered (3→4), and its success or failure is recorded. Regardless of success or failure, the gripper arm nudges the lid to reset its position (5). (B) Relationship between the offset from the origin and the probability of success (each n = 125).

**Table 2:** Operation sequence for evaluation of magnitude of positional misalignment of transfer microplates performed by Dobot M1 Pro.

| Step | Description |
| --- | --- |
| 1 | Lift the lid from the plate. |
| 2 | Move the offset distance ($\Delta x$ or $\Delta y$) horizontally. |
| 3 | Put the lid on the plate. |
| 4 | Initialize the position of the lid by pushing the edge of the lid. |

The distance of the artificially induced offset was determined as follows: First, as a preliminary experiment, we sought values that would result in a probability of success of approximately 0% and 100%, which were Δx = (1.80, 2.30 mm) and Δy = (2.10, 2.20 mm), respectively. Subsequently, evaluations were conducted at equal intervals of Δx = 0.1 mm and Δy = 0.05 mm within this range. We added a more detailed investigation of the y-direction because the results were only extremes (the probabilities of failure were 0.008, 0.007, and 1.000). We sequentially determined the measurement point as Δy = 2.175, 2.190, and 2.195 mm via binary search.

The operation of the robot arm was video recorded and analyzed using an in-house script (geometry_distribution.py) provided in our GitHub repository in Section 2.1. We used markers to determine the position of the microplate lid. For the markers, circular stickers with a diameter of 9.5 mm were used (Though-Spots, RNBW-1000, Diversified Biotech, Inc., MA, US) (Supplemental **Figure S3**). Initially, the RGB values of the stickers used as markers were obtained from the video data using OpenCV^27^. Thereafter, the coordinates of the observation point for determining the success or failure of the lid closing were determined, and success or failure was identified based on the distance between the center points of the yellow and blue stickers (Supplemental **Figures S3 B and D**). The recorded video is presented in Supplemental **Movie S1**, and the raw values of the determined successes or failures are presented in Supplemental **Table S2**.

## 3. RESULTS and DISCUSSION

### 3.1 Hardware setup for microplate transfer

We selected the Dobot M1 Pro as a selective compliance assembly robot arm (SCARA) robot^28^, within the category of multi-jointed robots, which excels in horizontal and vertical movements (**Figure 1A**). The action of placing microplates on the stage of a pipetting machine or retrieving them through horizontal and vertical movements is identical to a task referred to as "palletizing" in the factory automation industry. We chose SCARA robots because they are particularly suited to this task^28^ and have been adopted for automated solutions in biological experiments^29^. Many commercially available robotic gripper arms lack the width required to grasp microplates. Thus, we fabricated the fingers and mounted them onto the gripper using a 3D printer (**Figures 1B and C**). We constructed a hardware device to evaluate the accuracy of microplate transfer. The stage was created using a 3D printer, and the device was assembled using aluminum frames, digital gauges, and the microplate holder (**Figure 1D**). For detailed information on the device, please refer to Section 2.

### 3.2 Evaluation of position repeatability of transfer microplates

The position repeatability of microplate transfer was measured using dial gauges. We moved the microplate to a specific spatial coordinate using the Dobot M1 Pro, set the dial gauge reading at that point as (x, y) = (0, 0), and evaluated the position repeatability by repeating this movement multiple times based on the variation in the dial gauge readings.

Displacements were measured with the movement described in Section 2. **Figure 2A** presents an overview of the process employed to evaluate position repeatability. The microplate placed on the holder was grasped using the gripper and moved along the x and y directions to the measurement coordinate. Displacements in each direction were measured when the microplate body was pressed against the dial gauge’s probe. The microplate was then transferred to its previous position and placed on the holder. Thus, the microplate continuously traveled the path denoted as {(1) → (2) → (3) → (2) → (1)} in **Figure 2A**. This process was repeated 125 times at two transfer speeds: high

(100%) and low (50%), and the displacements in the x and y directions were measured (**Figure 2B**). As evident, the interquartile range (IQR) along the x and y directions under high-speed condition was 0.020 and 0.006 mm, respectively, and under low-speed condition it was 0.034 and 0.014 mm, respectively (raw data is presented in supplemental **Table S1**). The IQR along the x direction was larger than that along the y direction. The distribution in the x-direction was more pronounced than in the y-direction. This difference may be attributed to the orientation of the fingers holding the microplate.

### 3.3 Effect of magnitude of positional misalignment on microplate transfer success

We evaluated the effect of positional misalignment on the success of microplate transfer. After lifting the lid using a robot arm from the microplate, we artificially introduced a positional deviation of either Δx or Δy when placing it back on the microplate and created an operation sequence for such a transfer (**Figure 3A**). We repeated this transfer multiple times to evaluate the degree to which the probability of success decreased owing to the artificial positional deviation. The result of the transfer was judged a failure if the lid rode up on the microplate and could not return to its starting position. We added markers to the lid and counted the transfer successes by analyzing video images. Various distances of artificial offset for Δx or Δy (1.80–2.30 mm and 2.10– 2.20 mm, respectively) were introduced, and transfer was repeated 125 times for each case. The relationship between the distance of the offset and the probability of successful transfer is shown in **Figure 3B**. Along the x-direction, the probability of success gradually decreased with an increase in Δx. Whereas, along the y-direction, the probability significantly decreased when Δy exceeded 2.190 mm.

Next, we considered the physical shape of the circumference of the lid and microplate. We focused on the presence of a slope on the side of the microplate (**Figure 4**). Even if the lid collided with the microplate owing to the given misalignment, there may be cases wherein the lid slides down this slope and moves to the correct position. Such sliding occurs when the edge of the lid is even slightly engaged with the slope. Despite the thickness of the lid of approximately 1 mm and the slope on the side of the microplate spanning approximately 1 mm along the x-direction (**Figures 4C–E**), the misalignment distance was not contradicted when the probability of transfer failure began to increase, as shown in **Figure 3**. The different patterns of the probability of success change between x and y can be attributed to the differences in repetitive positioning accuracy, as shown in **Figure 2**. If the given positional misalignment was near the boundary (**Figure 4E**) that determines the success or failure of the transfer, poor repetitive positioning accuracy would result in variability in whether it exceeded the boundary owing to error. Conversely, with higher repetitive positioning accuracy, the variability in results can be expected to be minimized.

**Figure 4.**
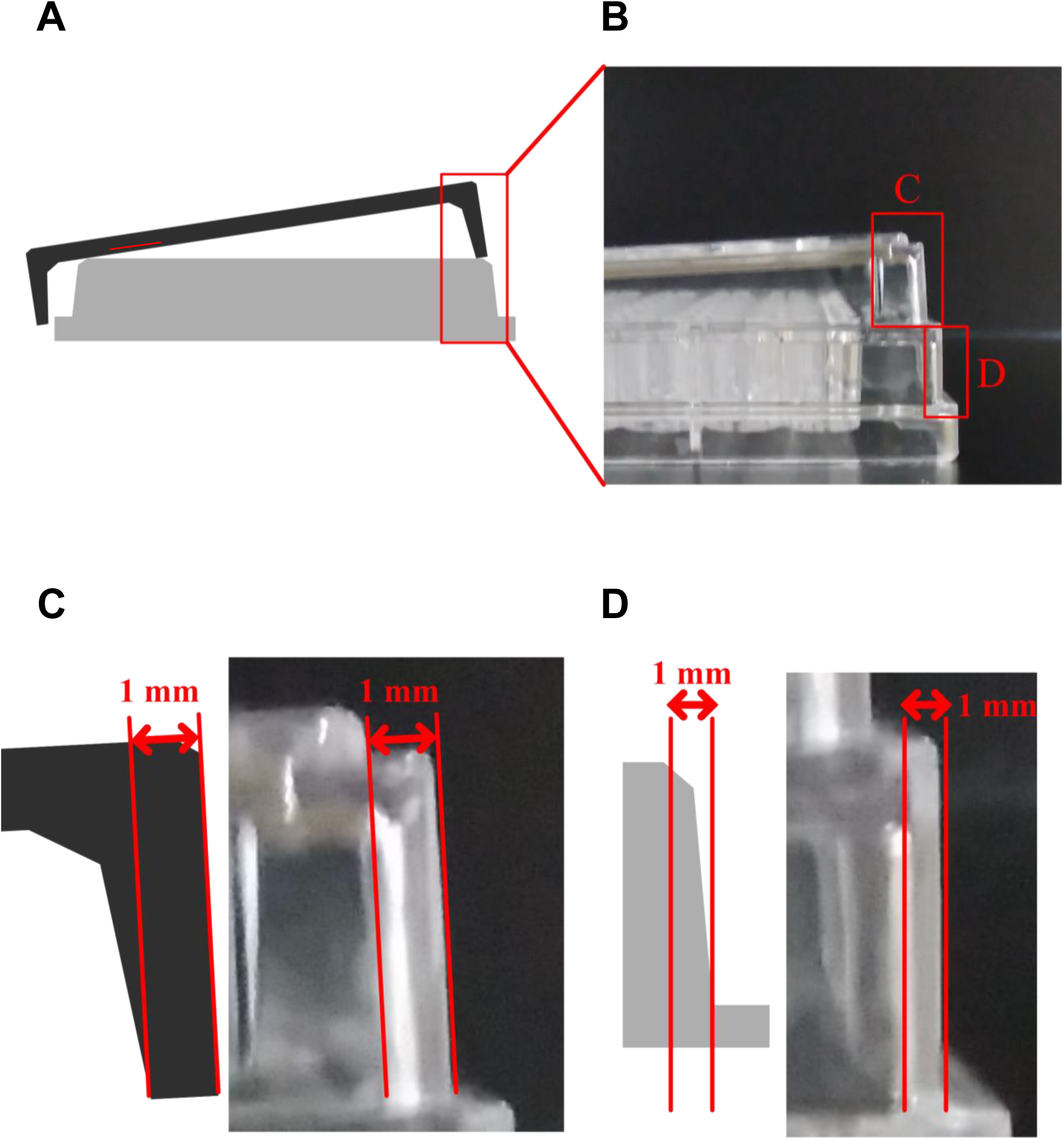
(A) Overview of the failure pattern of closing the lid of a microplate (corresponding to Figure 3A-4). (B) Photograph of a microplate and a misplaced lid. (C) Enlarged view of the lid. (D) Enlarged view of the microplate. (E) Enlarged view of the contact point between a microplate and the lid.

## 4. CONCLUSION

This study constructed a platform to verify the positional repeatability of a robotic arm when transferring microplates. We successfully evaluated its position repeatability using this platform. Moreover, by introducing artificial positional misalignments during the transfer of the lid of a microplate, we investigated their effect on the probability of successful transfers. These findings are expected to benefit biologists who are considering the adoption of robotic arms in terms of laboratory automation by assisting them in evaluating robot arms to be purchased. However, this study had certain limitations.

The results presented in this paper regarding positional repeatability and probability of successful transfer were specific to the hardware setup and parameters employed. Various factors can influence the positional repeatability and the probability of successful transport. For instance, the shape and material of the gripper and the labware being transferred can depend on rigidity and friction coefficients. The weight of the transferred items may be another crucial factor. Furthermore, the posture and the range of movement of the robot arm may have affected the results. To harness the robot arm more extensively in the field of biology, the effects of these conditions must be investigated in the future.

Our study did not consider experiments involving robots with different performances. We only tested Dobot M1 Pro; however, more options could be presented to biologists by testing other robots. Wider laboratory automation could be adopted if affordable robots such as the Dobot magician^16^ or myPalletizer^30^, priced at approximately 1000 USD, are demonstrated to be suitable for transfers. Although we only considered SCARA robots in this study, many industries utilize vertical articulated robots. SCARA robots are particularly suitable for palletization owing to their cost-effectiveness and high specifications. However, one of the benefits of vertical articulated robots is their flexible trajectory in space, which could potentially offer more stable transfer methods, such as accessing microplates diagonally^31^ to increase the probability of success during tasks like lid openings. The effect caused by the differences in the type of robot arm remains unknown and thus should be studied. Further, studies on different types of robot arms are necessary to utilize them for laboratory automation in biology.

In this study, we conducted experiments using an actual robot and microplates. Various tasks related to the mechanics of handling rigid bodies using robots can be addressed based on the research findings in the field of mechanical engineering. For instance, the movement of a rigid body on a chamfered slope, as shown in **Figure 4**, can be considered a sub-problem of a kinematically well-analyzed model system known as the peg-in-hole task^32^. Many computational and theoretical analyses of such sliding motions have been conducted. These insights can aid in understanding the influence of the differences in the features discussed above on robot transfers.

The platform constructed in this study has high reusability because of the adoption of an affordable and easy-to-handle dial gauge as a measuring device and the public 3D data for components such as the aluminum frame used to construct the platform. The layout, including the position of the measuring device and the stand for the microplates, can also be updated. These features can enable researchers to conduct further investigations and obtain results. In the future, the accumulation of investigations will deepen our understanding of the characteristics of tasks related to automation in biological experiments, and the introduction of robotic arms is expected to become easier. This could accelerate the proliferation of laboratory automation in biology.

## Acknowledgments

We thank the Laboratory Automation Supplier’s Association for their technical advice and thoughtful discussions. We thank Dr. Natsuki Yamanobe, Dr. Yasufumi Tanaka, and Mr. Kenji Matsukuma for the helpful discussion. We would like to thank Editage (https://www.editage.jp/) for English language editing.

## Supplemental materials

Figure S1.

Evaluation setting details.

Figure S2.

Fixing the aluminum flame stage on the experimental table.

Figure S3.

Appearance of the marker during evaluation of the success or failure of transfer when a positional deviation is introduced intentionally. (A) One of the flames in the original video. (B)–(D) Results of detection of each color (blue, green, and yellow, respectively). Red circle denotes the center of each sticker.

Table S1.

Preprocess data of positional repeatability measured by digital gazes.

Table S2.

Preprocess data of evaluation of the success or failure of transfer when a positional deviation is introduced intentionally.

Movie S1.

Video of the evaluation of the success or failure of transfer when a positional deviation is introduced intentionally.

https://drive.google.com/drive/folders/14_GV0-YAznDkDeLjCMLWuaTZydq-L2YC?usp=drive_link

File S1.

3D-CAD data of jigs created in this study.

File S2.

Size of each aluminum frame used to construct the experimental stage.

