## Supplementary figures and images for "Evaluation of Microplate Handling Accuracy for Applying Robotic Arms in Laboratory Automation"

### Supplemental Figures

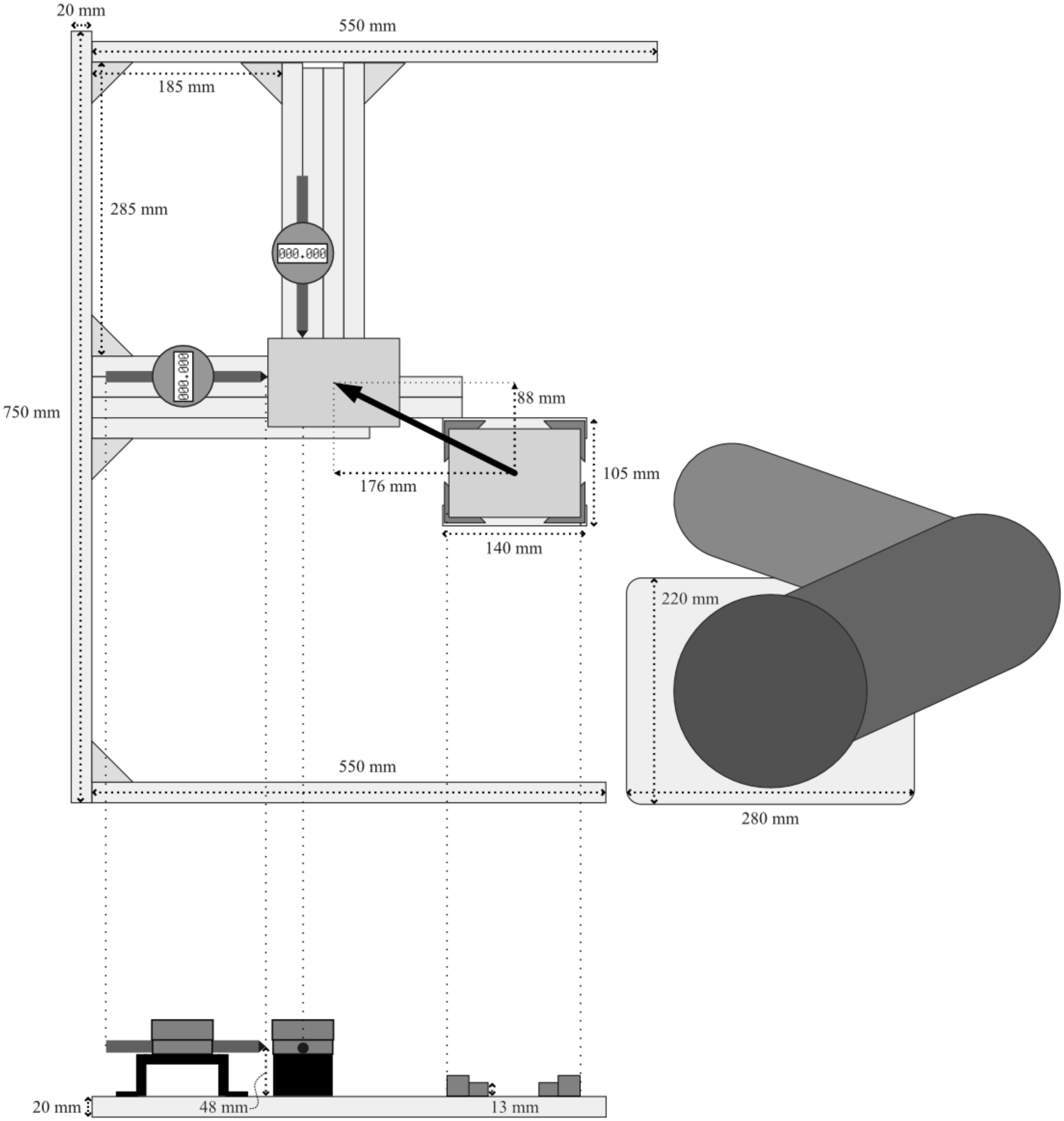


Harazono et al., Figure S1

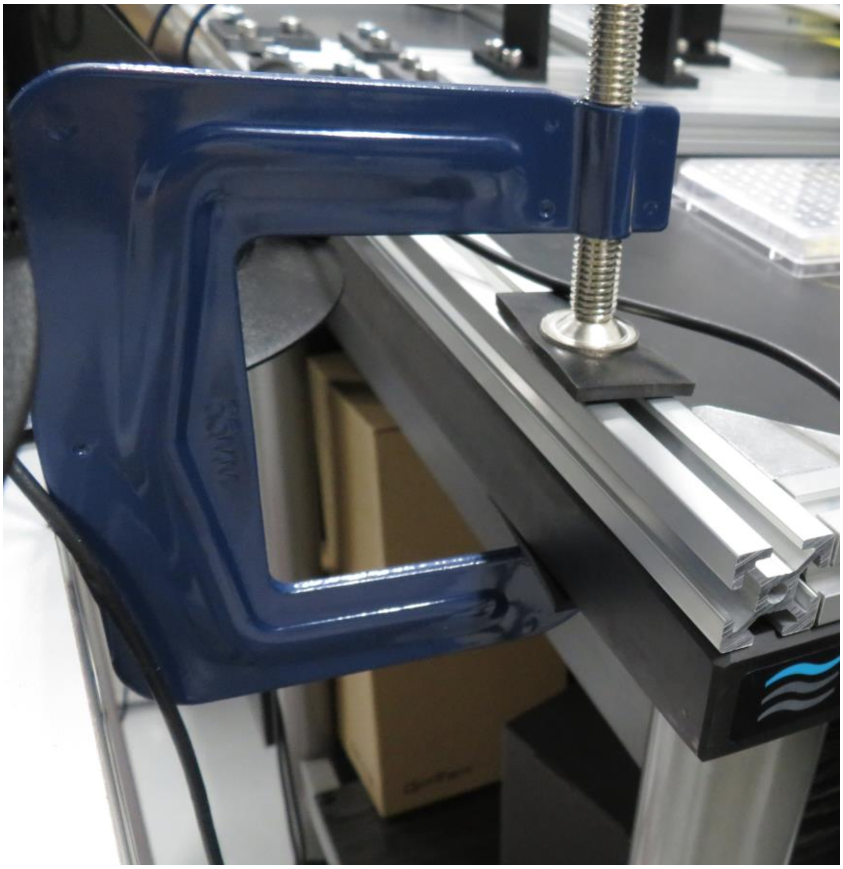

Harazono et al., Figure S2

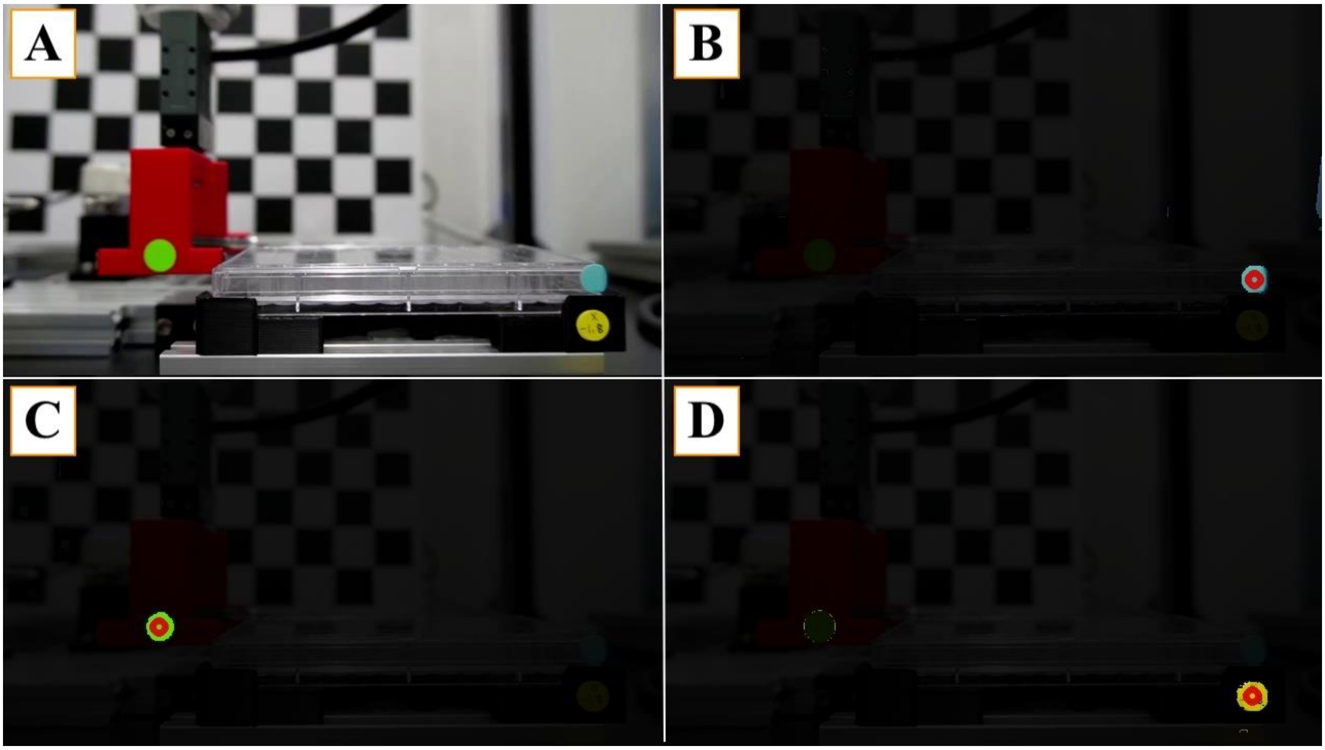


Harazono et al., Figure S3
