## Supplementary material for "Evaluation of Microplate Handling Accuracy for Applying Robotic Arms in Laboratory Automation": File S2: Size of each aluminum frame.pdf

| 見積ID | 組図ID | 型番 | 部品名 | 数量 |
|---|---|---|---|---|
| 1 | 1-1 | NFS5-2020-140 | 5シリーズ 正方形 20×20mm 1列溝 4面溝 | 1 |
| 2 | 2-1 | NFS5-2020-140-LTP | 5シリーズ 正方形 20×20mm 1列溝 4面溝 | 1 |
| 3 | 3-1 | NFS5-2020-270-LTP | 5シリーズ 正方形 20×20mm 1列溝 4面溝 | 1 |
| 4 | 4-1 | NFS5-2020-270-RTP | 5シリーズ 正方形 20×20mm 1列溝 4面溝 | 1 |
| 5 | 5-1~2 | NFS5-2020-280-RTP | 5シリーズ 正方形 20×20mm 1列溝 4面溝 | 2 |
| 6 | 6-1 | NFS5-2020-490 | 5シリーズ 正方形 20×20mm 1列溝 4面溝 | 1 |
| 7 | 7-1 | NFS5-2020-550 | 5シリーズ 正方形 20×20mm 1列溝 4面溝 | 1 |
| 8 | 8-1~2 | NFS5-2020-60 | 5シリーズ 正方形 20×20mm 1列溝 4面溝 | 2 |
| 9 | 9-1 | NFS5-2020-750 | 5シリーズ 正方形 20×20mm 1列溝 4面溝 | 1 |
| 10 | 10-1 | NFS5-2060-280-RTP | 5シリーズ 長方形 20×60mm 20×80mm 3列溝以上 4面溝 | 1 |
| 11 | 11-1 | NFS5-2060-360-TPW | 5シリーズ 長方形 20×60mm 20×80mm 3列溝以上 4面溝 | 1 |
| 12 | 12 | HBLFSN5-SET | 突起付反転ブラケット | 15 |

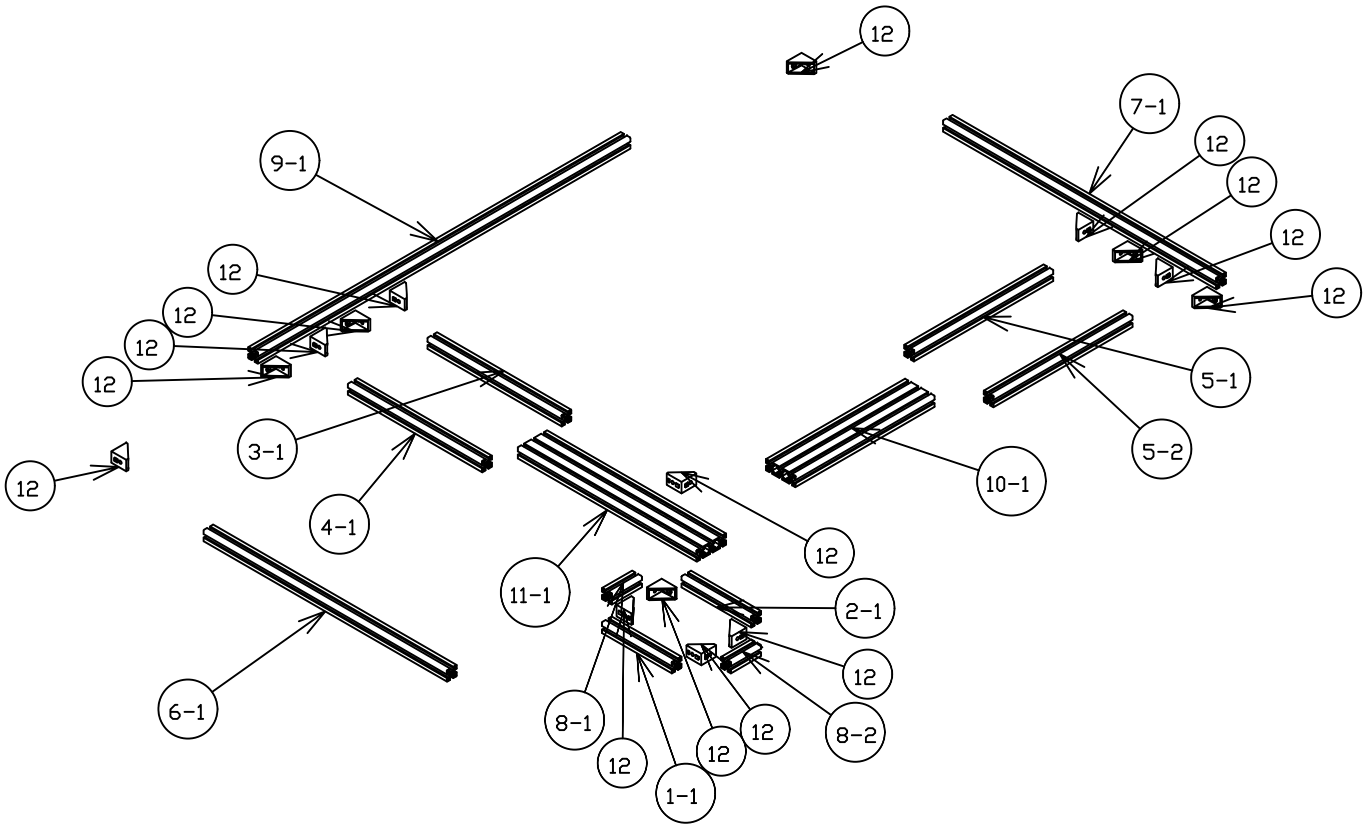
